# When things add up: environmental structure and microbial interactions drive antibiotic resistance plasmid evolution

**DOI:** 10.1101/2025.05.07.652652

**Authors:** Laurens E. Zandbergen, Mishelle Bustamante, Jan-Maarten Luursema, Thomas Hackl, Marjon G.J. de Vos

## Abstract

**Background and objectives:** Antimicrobial resistance is a major global health threat, driven in part by the rapid evolution of resistance in pathogens, which undermines the effectiveness of antimicrobial treatment. In infections, pathogens rarely live in a well-mixed, single species environment. It is an open question how microbial interactions in contrasting environmental structures affect plasmid mediated antibiotic resistance evolution.

**Methodology:** This study investigates how a spatially structured versus a well-mixed liquid environment, together with microbial interactions, affect antibiotic resistance evolution in uropathogenic *Escherichia coli.* We conducted a serial transfer experiment under increasing concentrations of trimethoprim-sulfamethoxazole comparing resistance evolution in the well-mixed and spatially structured environments, both in the presence and absence of a polymicrobial community.

**Results:** Our results revealed that *E. coli* in community context displayed parallel evolutionary trajectories, leading to higher final antibiotic tolerance, while the spatial structure allowed for prolonged resistance evolution. Copy number variation of the plasmid-borne resistance locus varied significantly across conditions; *E. coli* evolved in the well-mixed, monoculture conditions, exhibited the greatest increases in copy number, whereas lineages evolved in the presence of the community showed minimal changes relative to the ancestor.

**Conclusions and implications:** These findings underscore the complex interplay between the genetic basis of resistance, the environmental structure and microbial ecology in shaping plasmid-mediated antimicrobial resistance evolution.

**Lay summary:** Antibiotic resistance depends on environmental context, microbial interactions, and genetics. This study shows that *E. coli* evolved resistance differently in mixed versus structured environments, with changes in plasmid-borne resistance gene copy number. Community interactions led to similar evolutionary paths and higher tolerance, while well-mixed, single-species conditions drove larger genetic changes.

## Main Text

### Background and objectives

Polymicrobial communities can have a profound effect on the severity of infections^1–4^ and can diminish treatment efficacy of pathogens.^5–8^ Pathogens are increasingly becoming resistant to antimicrobials, complicating treatments.^9^ According to the World Health Organization, antimicrobial resistance (AMR) is among the top ten of threats to global health.^10^

Emerging research shows that AMR can be a collective trait that develops within bacterial communities.^7,11,12^ For example, tolerance to antibiotics can be increased in a community by decreasing antibiotic pressures in the local environment through antibiotic degradation, leading to increased pathogen density.^13–15^ Higher population densities have been shown to decrease the susceptibility to certain antibiotics, a phenomenon known as the inoculum effect.^16,17^ Additionally, microbial communities are often driven by competitive interactions, which lead to a reduction in population size per community member and therefore a change in survival.^18^ Yet, cooperation also occurs, particularly in stressful environments^19,20^, and it has been observed that interspecies interactions can lead to antibiotic protective effects.^11,21,22^

The environmental structure of bacterial communities can have a large impact on bacterial growth. Virtually all known bacteria are capable of forming biofilms or attaching to surfaces, which, through differences in lifestyle enhances their tolerance to environmental stressors, both biotic and abiotic.^23,24^ Surface attached spatial growth versus planktonic lifestyle in a well-mixed environment has different effects on the selective pressures and growth of bacterial populations, such as alterations in competitive and mutualism dynamics.^25,26^ Environmental structure has been implicated as a critical factor for the stability and coexistence of species in a community.^27–29^ For example, gene expression related to quorum sensing, and consequently antibiotic susceptibility,^22,30^ is different in a spatially structured environment compared to a well-mixed environment.^31^ Importantly, theorical and empirical research suggests that social interactions of bacteria in spatially structured environments are significantly altered compared to environments with better mixing, leading to different survival outcomes.^32–35^

Environmental structure and microbial lifestyle can influence ecological interactions among microbes. Such different ecological interactions can in turn affect microbial co-existence and evolution.^27–29^ Previous studies indicated that populations in spatially structured environments adapt more slowly than those in well-mixed environments ^25,36–38^, as evolution in a spatially structured environments increases genetic drift and founder effects in local populations reducing the rate of adaptation.^25,38^ Additionally, mixed environments typically lead to a decrease of metabolite exchange among species, focal species will therefore potentially be less affected by the presence of community members.^39,40^

Antimicrobial resistance is often conferred by plasmids containing antibiotic resistance genes. Microbial lifestyle, environmental and community context influence the emergence of plasmid encoded resistance^41–44^. It is unclear how these factors affect the repeatability of plasmid acquired antibiotic resistance evolution. We therefore investigate the effect of the presence and absence of a four-member microbial community of a focal pathogenic species, in two contrasting environments, a well-mixed liquid and a spatially structured solid-surface environment on the repeatability of antibiotic resistance evolution.

## Materials and methods

### Bacterial isolates and growth medium

The focal *Escherichia coli* isolate and three community isolates (*Klebsiella pneumoniae*, *Enterococcus faecium*, and *Staphylococcus epidermidis*) were collected anonymously in a previous study from elderly patients (>70 years old) who were suffering from a urinary tract infection.^45^ The three community members were picked based on their ability to grow in Lysogeny Broth (LB) containing 250 µg/mL trimethoprim and 1250 µg/mL sulfamethoxazole, and being able to clearly distinguish all four isolates on CHROMagar Orientation (CHROMagar). The *E. coli* isolate was selected based on its sensitivity to trimethoprim/sulfamethoxazole. Pure cultures of each separate isolate were obtained by streaking on CHROMagar plates, selecting a single colony, and growing overnight in LB broth at 37°C. Overnight cultures were used to create glycerol stocks (25% glycerol v/v) which were subsequently stored at -80°C. The experiments were performed in Artificial Urine Medium (AUM), which is created (for a 1X concentration) by mixing 1.5 g/L bacto peptone L37 (BD), 3.15 g/L NaHCO3, 11.25 g/L urea (Roth), 4.8 g/L Na2SO4.10H2O, 1.8 g/L K2HPO4, 1.95 g/L NH4Cl, 15 mg/L bacto yeast extract (BD), 1.98 mM lactic acid (Roth), 600 mg/L citric acid, 105 mg/L uric acid, 1.2 g/L creatinine, 4.44 mg/L CaCl2.2H2O, 1.8 g/L Fe(II)SO4.7H2O (Riedel), 368 mg/L MgSO4.7H2O, 1.43 g/L KH2PO4, 5.1 g/L NaCl. Chemicals were ordered from Sigma, unless stated otherwise. The pH was adjusted to 6.5 using 1M HCl.

### Antibiotic and X-gal stocks

The trimethoprim stock was created by mixing 10 mg/mL trimethoprim in dimethyl sulfoxide (DMSO), and the sulfamethoxazole stock was created by mixing 50 mg/mL sulfamethoxazole in acetone. The X-gal stock contained 20 mg/mL X-Gal and 0.1M Isopropyl β-D-1-thiogalactopyranoside (IPTG) mixed in DMSO. Both antibiotics, the X-gal, IPTG and solvents were ordered from Sigma. The trimethoprim stock was stored at 4°C for up to two weeks, and the X-gal stock and sulfamethoxazole stocks were stored at -20°C. Trimethoprim and sulfamethoxazole were always mixed in a 1:5 ratio, similar to clinical settings.^46^

### Bacterial hotel creation and transfer protocol

Silicon hotel agar molds were stored in 70% ethanol and were dried in a sterile flow cabinet before pouring agar. Each agar hotel was created by mixing 12.5 mL 2x AUM, an appropriate amount of antibiotics, 12.5 mL 3% soft agar in demineralized water at 60°C, and 125 µL X-Gal stock for the visualization of *E. coli* into the hotel mold. X-Gal makes *E.coli* turn blue. Agar hotels were cooled for 30 minutes before removing the agar from the molds.

Five replicates of *E. coli* were evolved for 60 bi-daily transfers in in isolation and in communities with *K. pneumoniae*, *E. faecium* and *S. epidermidis.* Starting cultures of all isolates were created by growing each separately in 1x AUM overnight at 37°C shaken at 200 rpm. Each culture was diluted a hundred times in 1x AUM. Starting agar hotels were created with the starting antibiotics concentration of 50:250 µg/mL TMP:SMX, this 1:5 ratio was picked to resemble the ratio used in clinical settings.^46^ Following the layout of Figure S4, 0.5 µL of each diluted culture was added to the middle of the appropriate well on the agar plate. Added cultures were allowed to dry for five minutes before incubating the agar hotels at 37°C for 48 hours. We did not observe any spillover of colonies to other compartments.

Before transferring *E. coli* to a fresh agar hotel, images of hotels were taken and based on the surface area of the *E.coli* colony, if the colony’s growth exceeded 1mm of the starting colony, the concentration of antibiotics would be increased by 20%. Unless the antibiotics concentration was increased during the previous transfer. Exception here were the first two transfers where antibiotics were increased by 20% due to clear overgrowth of *E. coli*. Colonies were transferred by streaking around the outer edges of each *E. coli* colony five times and suspending bacteria in 50 µL of 1x AUM. After mixing, 0.5 µL of each replicate would be added to the fresh agar hotels. Wild type community members (non-*E. coli*) were always added as hundred times diluted overnight culture in 1x AUM to the fresh agar hotels.

### Liquid serial transfers protocol

*E. coli* was evolved over 120 days in AUM transferring replicates to fresh 5 mL medium every 48 hours for 60 transfers in total. To start, the *E. coli* and the three community members were streaked on agar to obtain a pure culture of a single colony that was cultured overnight in AUM at 37°C, shaken at 200 rpm. Isolates were diluted a hundred times by adding 50 µL to 5mL total preheated medium. Five replicates of *E. coli* in isolation and five replicates of *E. coli* with the three community members *K. pneumoniae, E. faecium*, and *S. epidermidis*. An appropriate amount of the antibiotics were added to the medium before adding the bacteria. Antibiotic concentration started at 50:250 µg/mL TMP:SMX, 50% of the *E. coli* minimum inhibitory concentration. Cultures were allowed to grow for 48 hours at 37°C and shaken at 200 rpm. During transfer, the culture was diluted a hundred times and community members were added from stock. Remaining culture was used for obtaining CFU/mL counts for *E. coli* on CHROMagar plates. If *E. coli* had exceeded 1*10^6^ CFU/mL, antibiotic concentration would be increased by 20% for the next transfer. For the first two transfers for *E. coli* evolving in presence of the community, the antibiotic concentration was increased by 20% immediately due to observed high population growth based on turbidity.

### MIC and MBC assays

Minimal inhibitory concentrations (MIC) of replicates were measured in liquid. Using the concentration of antibiotics at the final transfer step as a guideline, a dilution series was created for each isolate specifically. Measurements were performed in 96-well microtiter plates, using 1 x AUM as growth medium. Stationary phase cultures of *E. coli* replicates were used for experiments, diluting them 40x times in the medium antibiotics’ mixture. Plates were incubated overnight at 37°C, shaken at 200 rpm. Growth was measured through absorbance at 600nm using a BMG CLARIOstar microplate reader. If the absorbance with the background subtracted exceeded 0.1, it was assumed that *E. coli* had grown using the lowest concentration of antibiotics where no growth was observed is the MIC. All MIC measurements were performed in triplicate.

Minimal bactericidal concentration (MBC) assays, which can be seen as extended MIC assays in which the community is also present, were performed following the above protocol, but with community members added together with the *E. coli* before overnight growth. After OD600 measurements to establish growth in the plate reader, the presence of *E. coli* was established by transferring 1µL using a 96-well pinning tool to CHROMagar, followed by overnight growth at 37°C. The lowest concentration of antibiotics where no growth of *E.coli* was observed on CHROMagar is the MBC.

### DNA extraction and sequencing

DNA from first transfer clones and final transfer mutants were extracted using the QIAamp DNA Mini Kit (Qiagen, Hilden, Germany) following the manufacturer’s instructions. DNA was eluted in 100 µL elution buffer AE. Quality and quantity of isolated DNA were checked using a Nanodrop One^C^ Microvolume UV-Vis Spectrophotometer (Thermo Fisher Scientific, USA).

Three clones of the final transfer step for each of the five replicate lineages per experimental setting were sent for illumina sequencing at SeqCenter (Pittsburgh, PA, USA). Five replicates of community members isolated from *E. coli* community agar and liquid were also sent to SeqCenter, Pittsburgh, USA for Illumina sequencing.

Illumina sequencing libraries were prepared using the tagmentation-based and PCR-based Illumina DNA Prep kit and custom IDT 10bp unique dual indices (UDI) with a target insert size of 320 bp. No additional DNA fragmentation or size selection steps were performed. Illumina sequencing was performed on an Illumina NovaSeq 6000 sequencer in one or more multiplexed shared-flow-cell runs, producing 2×151bp paired-end reads. Demultiplexing, quality control and adapter trimming was performed with bcl-convert1 (v4.1.5). In total 10.9 Gbp and 359.2 Gbp of read data were produced from ancestral and evolved strains, respectively.

Ancestor strains were sent to the same company for Nanopore sequencing. Sample libraries were prepared using Oxford Nanopore Technologies (ONT) Ligation Sequencing Kit (SQK-NBD114.24) with NEBNext® Companion Module (E7180L) to manufacturer’s specifications. All samples were run on Nanopore R10.4.1 flow cells and MinION Mk1B device. Post-sequencing, Guppy (v6.3.8) was used for super high accuracy base calling (SUP) and demultiplexing. Overall, 2.9 Gbp of read data were produced.

### Genome assembly, annotation variant calling and effect prediction

The wildtype reference assembly was generated with flye v2.9.2 (settings: --nano-hq)^47^ from the ancestral nanopore data and polished using ancestral Illumina read with polca.sh included in the MaSuRCA v4.1.0 Genome Assembly and Analysis Toolkit.^48^ Genes were annotated with Bakta v1.8.1^49^ and integrated prophages and plasmids with genomad v1.7.4^50^ for each replicate were called from read mappings generated with minimap2 v2.26^51^ using freebayes v1.3.1 (settings: -p 1 -0 -F 0.5)^52^ and filtered for spurious calls with bcftools v1.17 (--exclude ‘REF∼“N” || QUAL < 20 || INFO/DP < 10’).^53^ Variant effects were predicted with the Ensembl Variant Effect Predictor (VEP) v110.1 command line tool.

### *H*-index calculation

To quantify the repeatability of mutations within our *E. coli* mutants, the *H*-index of genome repeatability previously described by Schenk *et al*. was calculated.^54^ The *H*-index was calculated on the functional (COG) and at the gene level. The final *H*-index value falls between 0 (no repeated evolution) and 1 (complete parallelism). An *H*-index value of 0.5 indicates that two compared mutants have half of their mutations shared between them.

## Statistical analysis

When comparing the values between two conditions of mutants, an ANOVA and two-tailed *Welch’s t*-test were performed to test the null hypothesis that the two parameters do not have significantly different values (hypothesis of neutrality). To determine level of correlation between two variables, we used the square of a Pearsson correlation coefficient and resulting *P*-value where the null-hypothesis was rejected with a significance level above 0.05. For multiple pairwise comparisons, we used independent sample *t*-tests for all sets with a Bonferroni-correction to adjust *P*-values to reduce type I errors.

## Results

### Environmental structure and microbial community impact evolutionary trajectories

To study the effects of structure and community context on the evolution of antibiotic resistance, we evolved a uropathogenic isolate of *E. coli* carrying a plasmid encoding resistance genes that only confer low resistance in a serial transfer evolution experiment under four different conditions. Evolving cultures were transferred bi-daily over 120 days, in artificial urine medium (AUM) containing increasing levels of trimethoprim-sulfamethoxazole. In each condition five replicate cultures of *E. coli* were evolved in both a (i) well-mixed and (ii) spatially structured AUM environment, both in the (iii) presence and (iv) absence of a community of three other bacterial species. These other species are often co-isolated with *E. coli*; *K. pneumoniae, E. faecium* and *S. epidermidis* in polymicrobial urinary tract infections (Materials and Methods).^45^

Phenotypic evolutionary trajectories of *E. coli* across replicate lineages overlapped for three of the four conditions. Replicate lineages overlapped for *E. coli* evolved in the spatially-structured environments in the presence and absence of the community. Replicate lineages of *E. coli* evolved in the well-mixed environment in the presence of the community also overlapped (Figure 1AB), but not in the absence of the community. Thus, the presence of the community leads to the canalization of the phenotypic evolutionary trajectories of *E. coli* in the well-mixed environment (Figure 1B).

**Figure 1.**
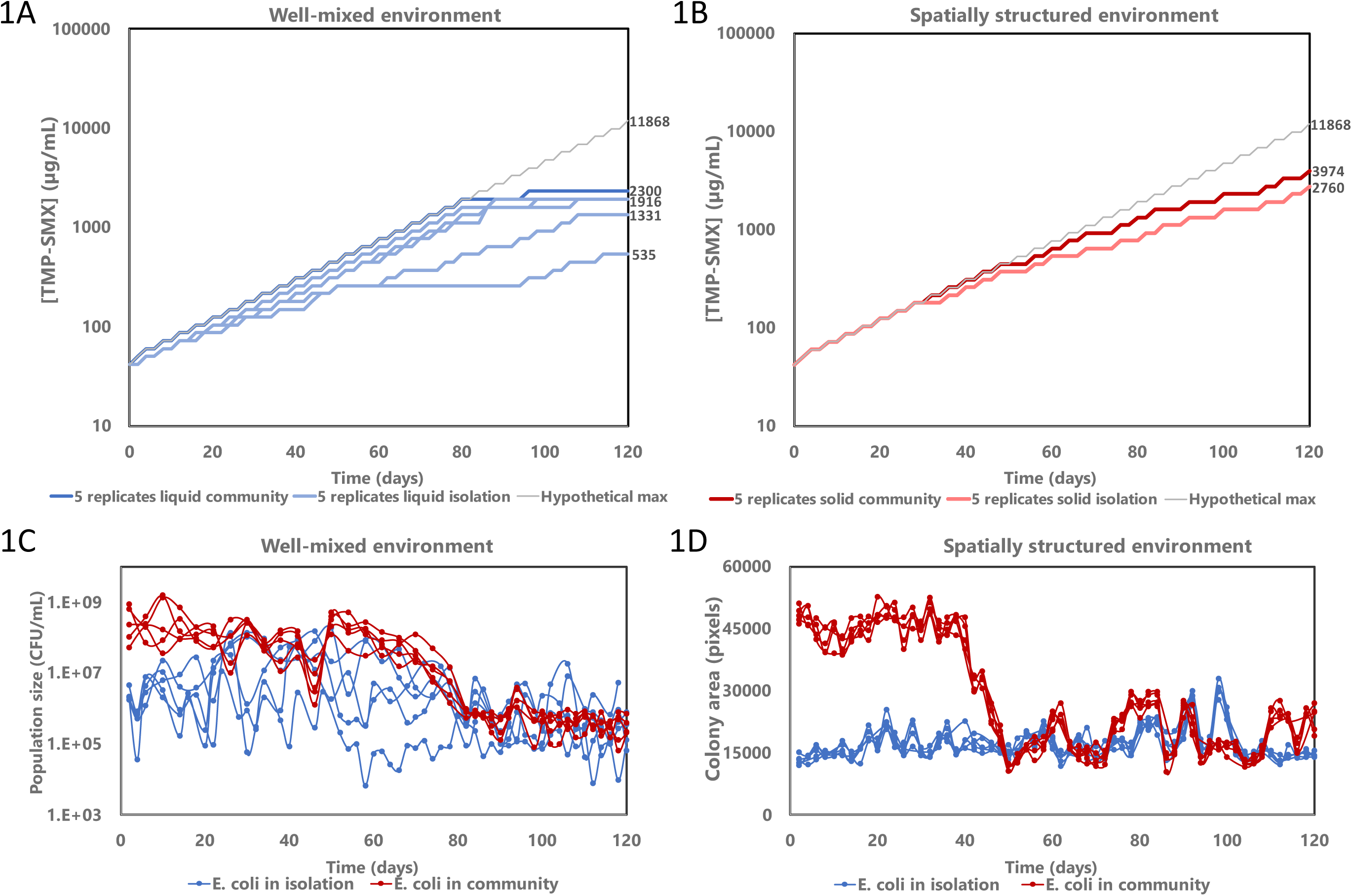
Phentoypic trajectories of trimethoprim-sulfamethoxazole antibiotic resistance evolution and population sizes of *E. coli* over time. *E. coli* evolved in the presence and absence of a community, consisting of *Klebsiella pneumoniae*, *Enterococcus faecium* and *Staphylococcus epidermidis*. *E. coli* was evolved in presence of trimethoprim-sulfamethoxazole (TMP-SMX) at a 1:5 ratio. Each condition was replicated five times. **AB,** concentration of TMP-SMX in the medium for all evolving replicate lineages within the well-mixed liquid environment (A) and spatially structured environment (B) over time. Concentrations of TMP-SMX are given by the trimethoprim concentrations. All five replicate lineages converged and overlapped for *E. coli* evolving in the presence of the community, for both well-mixed spatially structured environments. Different trajectories can be observed for *E. coli* evolving in the absence of the community within the well-mixed liquid environment. **C,** population size of *E. coli* in the well-mixed liquid environment in CFU/mL. A decline of the *E. coli* population size evolved in the presence of the community can be observed after forty eight days, for all five replicate lineages. The population size of *E. coli* evolving in isolation showed large fluctuations in the first eighty days, these became less diverse after this time. **D,** population size of *E. coli* evolved in the spatially structured solid environment measured as colony area in pixels. A clear decline of colony size for *E. coli* evolved in the presence of the community can be observed after roughly forty days for all five replicate lineages.

For the three overlapping replicate trajectories, the rate of evolution in initial weeks approached the hypothetical maximum rate. For the well-mixed environment, this persisted until day 80, whereas in the spatially structured environment the maximum rate lasted until day 48 for the *E. coli* evolved in presence of the community and until day 30 for *E. coli* evolved in isolation (Figure 1AB). These patterns suggests that within each of these environments, selective pressures drove evolution in a specific direction, potentially funneling evolution towards similar evolutionary outcomes and resulting in convergence of the phenotypic trajectories.

The presence of the community increased the level of antibiotics that *E. coli* can tolerate in both the well-mixed and spatially structured environments (*P*=0.023 for the community impact in the well-mixed environment and *P*<0.001 for the community impact in the spatially structured environment, *Welch’s t-test*). The final concentration of antibiotics that the evolved *E. coli* populations could sustain were higher in the community context than in isolation in both environments.

In addition to community context, environmental structure, either well-mixed liquid AUM versus spatially structured solid AUM-agar, influenced the evolutionary trajectories of *E. coli*. Initially, the rate of antibiotic increase was identical for both environments. However, after 48 days of evolution (24 transfers) the trajectories began to diverge. In the spatially structured community environment, the rate at which the antibiotic could be increased declined slightly, compared to the well-mixed community environment. Despite this slower rate, *E.coli* evolving at the spatially structured environments tolerated higher concentrations of antibiotics compared to the well-mixed conditions at the final transfer (Figure 1AB). We note that the antibiotic solvents could have influenced adaptation at the end of the experiments when antibiotic concentrations are high. Overall, these results demonstrate that environmental structure differentiated the course of the evolutionary trajectories, with well-mixed liquid and solid-surface environments following distinct evolutionary paths (one-way ANOVA, *P<*0.001 for liquid versus solid). These results show that both community presence, as well as environment structure had an impact on the evolutionary trajectories of trimethoprim-sulfamethoxazole resistance of *E. coli*. Community context promoted parallelism among evolutionary trajectories and higher final antibiotic tolerance, while spatial structure allowed for prolonged resistance evolution and higher final antibiotic concentrations than those in well-mixed environments.

### Community interactions mediate response to selection

Uropathogenic *E. coli* is known for its flexible metabolism, enabling it to adapt and respond to fluctuating nutrients in the environment.^55^ We investigated whether *E. coli* would respond differently to the presence of antibiotics in a community context. We observed that the focal *E. coli* outgrew the other community members. During the initial 30 days the density of *E. coli* was between the 10^8^ and 10^9^ CFU/mL for most replicates at the time of transfer in the well-mixed environment in the presence of the community, compared to between 10^6^ and 10^7^ CFU/mL in isolation (Figure 1C, *P*<0.001 for the first 30 days, ANOVA). This high population density in the presence of the community was maintained even after the antibiotic concentration reached the initial MIC at day 12. Even though the community members are highly resistant (>1:5 mg/mL TMP:SMX), *E. coli* achieved higher population densities from the onset of the experiment.

Similarly, in the spatially structured environment, *E. coli* benefited from the presence of the community; colonies surrounded by the community members were approximately three-fold larger in surface area compared to *E. coli* colonies surrounded only by other *E. coli* colonies (Figure 1D, *P*<0.001 for the difference between community and isolation colonies, *Welch’s t-test*). Population sizes among the five replicates lineages evolved within the same environment did not differ significantly (Figure 1CD, *P*>0.05, independent samples *t*-test with Bonferroni correction).

In the well-mixed liquid environment, the presence of the three-species community accelerated the rate at which the antibiotic could be increased in the experiment. When *E.coli* was grown in the presence of the community, *E. coli* lineages surpassed the wild type MIC level after 12 days, whereas lineages evolved in isolation required 16-18 days (*P*<0.001, ANOVA). In contrast, in the spatially structured environment, all replicates reached the native MIC after 12 days, regardless of whether *E.coli* evolved in the presence or absence of the community and despite the larger colony sizes of *E.coli* in the presence of the community.

The beneficial effects of the community on *E.coli* population size diminished over time. In the well-mixed environment, the rate of antibiotic increase slowed down after approximately 80 days (Figure 1A, dark blue graph). This slow-down occurred simultaneously across all five replicate lineages, which suggests that the community canalizes phenotypic trajectories in a consistent manner. Such a synchronization of the phenotypic evolutionary trajectories was not observed for the replicate lineages of *E. coli* in isolation (Figure 1A, light blue graphs). Some of the replicate lineages of *E.coli* evolving in isolation leveled off much earlier in the experiment, showing a slower phenotypic evolutionary trajectory, with a lower final level of antibiotics. For the spatially structured environment, such a pronounced slow-down and diminishing effect of the community were not observed. The antibiotic concentrations reached at the final transfer were higher for all lineages evolved in the spatially structured solid environment, than for those in the well-mixed environment, both in the presence and absence of the community (Figure 1B).

Also the population sizes of the community members declined over time. *K. pneumoniae* was initially abundantly present in the well-mixed environment (between 10^7^ and 10^8^ CFU/mL), but steadily decreased to between 10^6^ and 10^7^ CFU/mL after 14 days. At later time points, *K. pneumoniae* was no longer detected, indicating an abundance at least 100-fold lower than that of *E. coli*, which was below the detection limit. In the spatially structured environment, *K. pneumoniae*, steadily decreased in colony size during the first 14 days, consistent with population dynamics in the well-mixed environment. Colony sizes continued to decline with every increase in antibiotic concentration, indicating that rising antibiotic levels affects the wild-type *K. pneumoniae* fitness negatively (average colony size at day 2, 14494 pixels, at day 120, 8518 pixels).

*E. faecium* was initially present in low numbers (around 10^2^ -10^3^ CFU/mL), as expected based on previous studies,^4,11^ yet population sizes declined with increasing antibiotic concentrations, decreasing by at least five-fold between over the 120 day experiment. *S. epidermidis* was rarely detected during the experiments, indicating population sizes below 10^3^ CFU/mL, even though *S. epidermidis* is typically more abundant in isolation (10^4^-10^5^ CFU/mL). This suggests that competition with other species, or the increasing antibiotics concentrations limited the growth of *S. epidermidis*, even though it can survive high antibiotic concentrations^8^. The decline of both the focal species and community members over time could point to increased competition, reduced cooperation, or a general fitness decrease associated with the rising antibiotic concentration in the environment.

### Environmental structure and community effect on evolved antibiotic resistance level

Comparing the resistance level of *E. coli* lineages evolved in isolation with the lineages evolved in the presence of the community, we found that the lineages evolved in the well-mixed environment and the spatially structured environment differed. Although lineages evolved in the spatially structured environment tolerated higher concentrations of antibiotics at the end of the experiment than those in the well-mixed environment (Figure 1AB), we find that lineages evolved in the spatially structured environment exhibited lower MICs than those in the well-mixed environment when evolved in the presence of the community (resp. 0.8:4 mg/mL TMP:SMX and 3.2:16 mg/mL TMP:SMX, Figure 2A, Materials and Methods). This discrepancy suggests that different lifestyle (surface or liquid growth) or population dynamics affected antibiotic resistance evolution.

**Figure 2.**
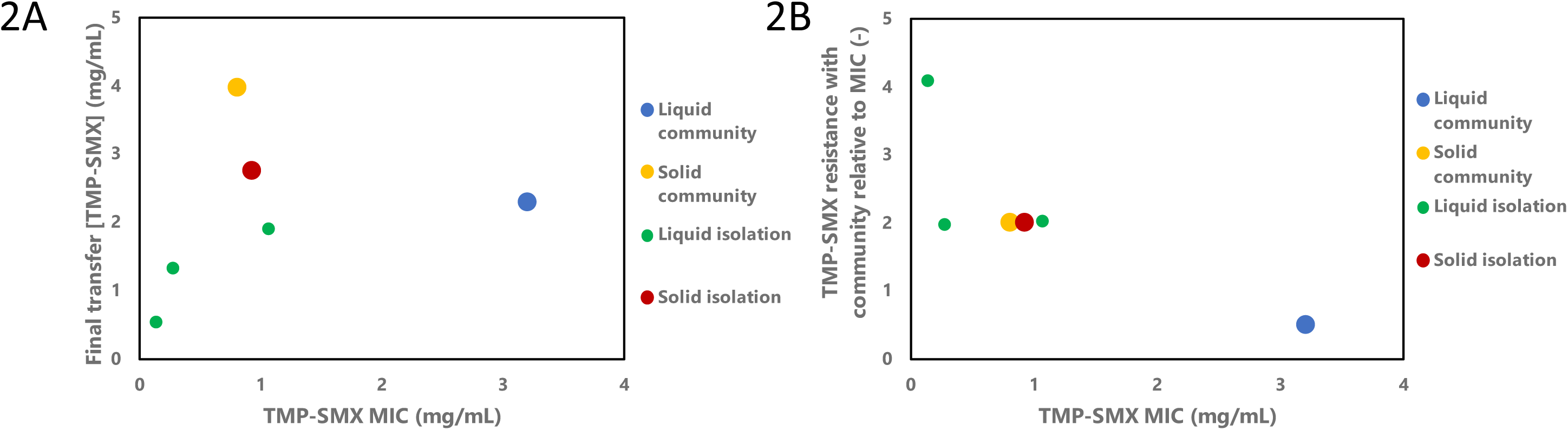
Relationship between the minimal inhibitory concentration (MIC) of evolved clones, in the presence and absence of the community, and the antibiotic concentrations at final transfer. The size of the data points shows the number of replicates, with the smallest data point being one replicate and the largest all five replicates. *E. coli* was evolved in presence of trimethoprim-sulfamethoxazole (TMP-SMX) at a 1:5 ratio. Concentrations of TMP-SMX are given by the trimethoprim concentrations. **A,** MIC of evolved *E.coli* versus the concentration of antibiotic at the final transfer step at day 120. No clear correlation between the MIC of evolved clones, and the concentration of antibiotic in the medium at their final transfer could be observed. **B,** MIC of evolved *E.coli* versus the community effect on MIC. The community resistance effect was determined by performing an MBC assay in the presence of the community, followed by subsequent transfer on agar. The concentration of trimethoprim-sulfamethoxazole at which no *E. coli* could be observed on the agar, is given relative to the MBC of *E. coli* in isolation. Therefore, a value above 1 on the y-axis indicates a higher observed resistance compared to measured MIC, while a value below 1 indicates a low resistance compared to the MIC. Only *E. coli* evolved in liquid medium in the presence of the community did not benefit from the presence of the community in the community MBC assay (blue). Absolute resistance levels in the presence of community are depicted in Figure S1.

A similar pattern was observed for *E. coli* evolved in isolation in both environments. In the spatially structured environment, lineages tolerated higher antibiotic concentrations at the final transfer compared to those in the well-mixed environment. However, the MICs of *E. coli* evolved in the spatially structure environment were lower (MIC 0.92:4.6 mg/mL TMP:SMX) than those of three of the five replicates from the well-mixed environment, (MIC 1.06:5.3 mg/mL TMP:SMX; *P*<0.001, *Welch’s t-test*). The remaining two replicates in the well-mixed environment had a lower MIC (0.27:1.33 and 0.13:0.66 mg/ml TMP:SMX) and corresponded to those lineages that sustained a lower final antibiotic concentration in the evolution experiment (Figure 1A, light blue graphs, Figure 2A).

Based on the observation that *E.coli* population sizes increased in the presence of community members, we hypothesized that *E. coli* benefited from one or more of the community members during the evolution experiment. To investigate this, we measured the minimum bactericidal concentration (MBC) at which the focal species could no longer be detected when cultured in the presence of the community (Figure 2B, Figure S1, Materials and Methods). We found that *E.coli* lineages evolved in the well-mixed environment in absence of the community could sustain twice the concentration of antibiotics when cultured in the presence of the community (MBC of 1.6:8 mg/mL TMP:SMX for 3 out of 5 replicates), compared to their MIC in isolation (0.8:4 mg/mL TMP:SMX for those same replicates) (Figure S1). In contrast, for lineages evolved in the well-mixed environment in the *presence* of the community, we observed a lower MBC compared to the MIC in presence of the community (MIC of 3.2:16 mg/mL TMP:SMX, MBC 1.6:8 mg/mL TMP:SMX). Indicating that the community negatively affected the growth of *E. coli* lineages that evolved in the presence of that community in the well-mixed environment. Interestingly, *E. coli* lineages evolved in the spatially structured environment benefited from the presence of the community; sustaining a two-fold higher concentration compared to the MIC (Figure 2B). These results indicate that the community generally has a positive effect on antibiotic tolerance of *E. coli* even after resistance evolution, except when *E. coli* evolved in the presence of the community in the well-mixed environment.

### Copy number variation of resistance loci on plasmid

To gain insights into the genetic basis of the evolved antibiotic resistance, we sequenced three clones from the final transfer of each of the five replicate lineages evolved under each condition and compared these sequences to that of the ancestor (Materials and Methods, Table S1). The ancestral strain, despite being highly sensitive to the start concentration of TMP-SMX at the beginning of the experiment, did carry a 138K base pair sized IncFII-plasmid that contains genes annotated to encode for resistance against both antibiotics, *dfrA17* and *sul1*. *DfrA17* confers resistance to trimethoprim, s*ul1* encodes a dihydropteroate synthase, conferring resistance to sulfamethoxazole. These resistance genes are located in an Tn21-like integron cassette, which is similar to a cassette first described by Liebert in 1999 (Figure S2).^56^ At the time of the description, the transposon did not yet contain the *dfrA17* gene, or any of the other resistance genes located on that cassette.

We assessed the plasmid copy numbers in the evolved clones and in the ancestor. The *E. coli* ancestor contained two copies of the plasmid relative to the host chromosome (Figure 3A). The evolved lineages had a similar plasmid copy number, indicating that plasmid copy number could not explain the increased resistance level. However, we observed copy number variations in the integron cassette containing the antibiotic resistance genes (Figure 3A). Zooming in on the resistance locus where the *dfrA17* and *sul1* genes are located, we observed substantial CNVs across conditions, replicates and even among clones from the same replicate lineage (Figure 3A, Figure S3). Overall, resistance locus CNVs were highest in clones evolved in isolation, in the well-mixed environment (median 5.2) and in the spatially structured environment (median 3.4). In contrast, copy numbers in *E. coli* evolved in community context were seemingly unaffected across the majority of clones (median 1.7 in the well-mixed and 2.0 in the spatially structured environment). Some clones deviated from the trend (Figure S3). The copy number variations (CNVs) of this cassette do not correlate with the MIC (*R*^2^=0.12, *P*=0.13, Figure 3B). Some lineages evolved in isolation in the well-mixed environment had a relatively low MIC despite having higher copy numbers of the antibiotic resistance genes (Figure 3B). This suggests that the MIC is not only influenced by resistance gene copy numbers, but also by other factors.

**Figure 3.**
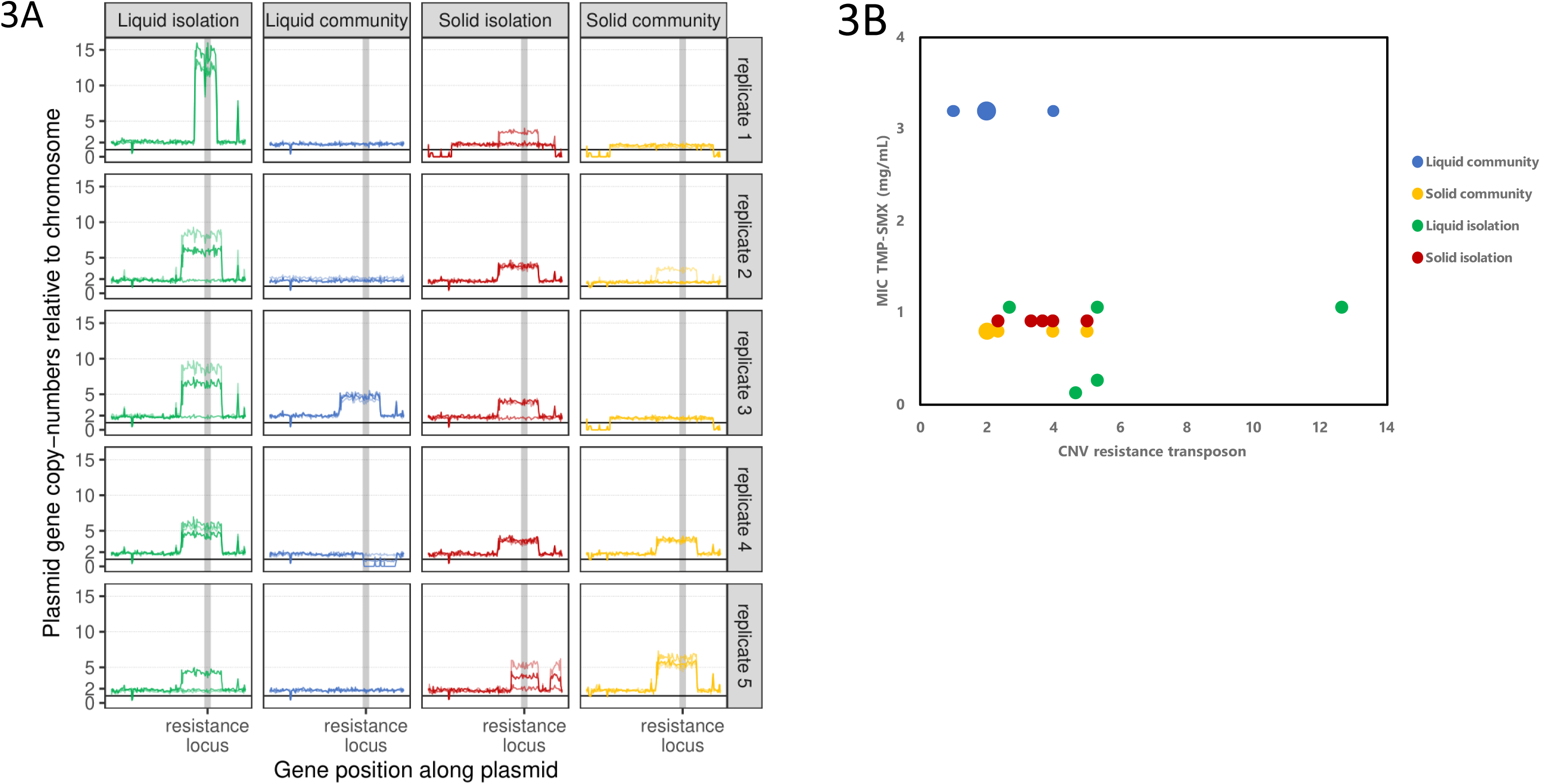
Copy number variation in the plasmid containing the antibiotic resistance genes in the final transfer mutants. Three clones were sequenced per replicate, showing genetic copy number variation within the population. **A.** Copy number variation in the TN21-like integron cassette part of the plasmid containing *dfrA17* and *sul1,* which confer resistance to trimethoprim and sulfamethoxazole, respectively, in each of the sequenced clones. The highlighted region in gray contains the resistance genes, showing overall a tendency for increased copy numbers of this region. Some replicates do not have increased copy numbers of the resistance genes compared to the wild type, particularly those evolved in the liquid environment in the presence of the community (blue), and in the solid spatially structured environment in the presence of the community (yellow). **B.** Average copy number variation per replicate in the TN21-like integron cassette part of the plasmid containing *dfrA17* and *sul1* resistance genes versus resistance level (MIC) of evolved replicate lineages. Concentrations of trimethoprim-sulfamethoxazole (1:5 ratio) are given by the trimethoprim concentrations. No clear pattern between replicates evolved in the same environment could be observed. Copy number variation for each sequenced clone is depicted in Figure S3.

Statistical analysis confirmed that in each evolutionary context the resistance locus CNV differed significantly from the CNV in the well-mixed environment in isolation (Figure S3). This shows that both community presence and environmental structure during evolution influence CNV of antibiotic resistance loci.

The implications of these data are multifaceted: overall, we found that lineages evolving antibiotic resistance in isolation had higher copy numbers of the *dfrA17-sul1* resistance loci, suggesting selection for increased gene copy numbers during resistance evolution under these conditions. In contrast, lineages evolving in the community context, showed no CNV increase in the majority of replicates and clones (7 out of 10 replicates or 20 out of 30 clones), with only moderate copy number increases in three replicates, indicating a distinct selection regime in the presence of the community. We noted previously that the presence of the community positively affects the growth of *E. coli*. We therefore hypothesize that this community-mediated growth effect increases antibiotic tolerance, and, consequently reduces selection for higher copy numbers of resistance genes for *E. coli* evolved in the presence of the community^57^.

### Environmental and community context modulate genetic adaptation

The number of genomic mutations within the evolved *E. coli* genomes differed significantly between environments. In the well-mixed liquid environment, we detected 25 mutations in total across all clones evolved in isolation, and only 16 mutations in clones evolved in the presence of the community. For the spatially structured environment, we detected in total 45 mutations in clones evolved in isolation, and 48 in clones evolved in the community context (*P*=0.004, *Welch’s t-*test comparing the well-mixed and the spatially structured environments). These findings align with our expectations, as a spatially structured environment versus a well-mixed environment is typically associated with diminished selection and increased genetic drift within the population,^58,59^ and a lower chance of beneficial mutations sweeping through the population.

Because many replicates evolved in the same environment followed parallel phenotypic trajectories (Figure 1AB), we hypothesized that lineages evolving under similar conditions would acquire similar genetic substitutions. To quantify genetic parallelism we calculated the *H-index*,^54^ which takes into account the fraction of shared mutations between two isolates on both the gene-level and the Cluster of Orthologous Group (COG) level for substitutions in the *E. coli* chromosome (Figure 4, Materials and Methods). In other work, we have observed quite a high degree of genetic parallelism in replicate populations evolved under the influence of TMP-SMX and bacterial interactions.^15^ In contrast, in this study, aside from the CNVs of the resistance genes on the Tn21-like integron cassette, we observed little genomic parallelism, with the exception of replicates evolved in the same spatially structured environment (Figure 4A). This indicates that, in the well-mixed environment, genomic mutations were mostly unique to individual replicates (Table S1). An exception involved two *E. coli* replicates evolved in the well-mixed environment in the presence of the community, which shared mutations in one gene. This mutated gene, *folP*, a known target of sulfamethoxazole ^46^, would allow *E. coli* to tolerate higher antibiotic pressures.^60^ The mutations observed, with the exception of *folP*, are, as far as we are aware of, not directly linked to antibiotic resistance to TMP-SMX.^54,60,62^ This mutation, in conjunction with other mutations in these replicates, did not result in significant population size differences compared with the other three replicates evolved within the same environment (*P*>0.05, independent samples *t-*test with Bonferroni correction). We did not identify any other chromosomal mutations in genes known to confer resistance in the evolved populations (Table S1).

**Figure 4.**
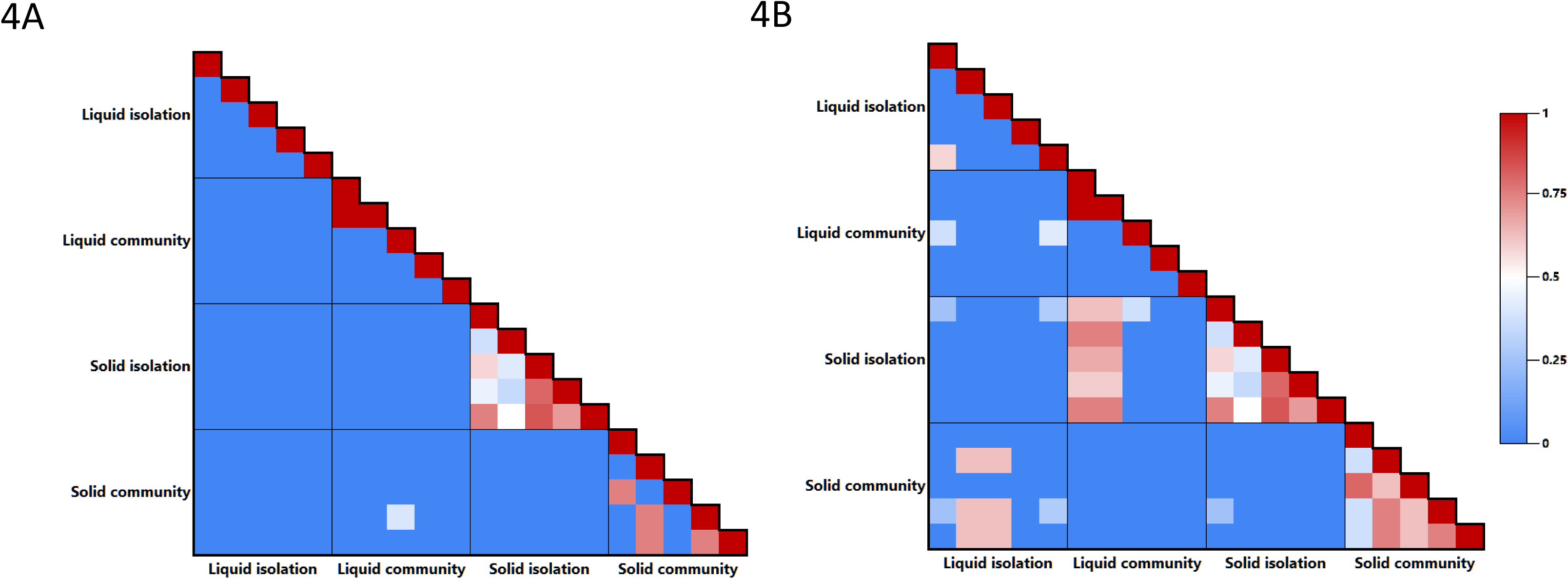
Clustering and mutation repeatability of final replicates clustered on gene and COG level. The level of genetic parallelism in evolved clones is given by the *H-*index. Higher *H-*index values indicate a higher parallelism of the two genotypes compared. Three clones per replicate were sequenced per environment. Comparisons in the blocks around the diagonal in each figure show the intra-environment comparisons, while the other blocks show the inter-environment comparisons. Solid indicates the spatially structured environment and liquid indicates the well-mixed environment. Mutations and genes are shown in Supplementary table 1. **A,** *H*-index of mutation repeatability at the gene-level of *E. coli* mutants. Within the spatially structured environment there was some repeated evolution, while very few replicates within the well-mixed liquid environment showed repeated evolution. **B,** *H-*index of mutation repeatability at the COG-level. Most genetic parallelism is found in the spatially-structured environment.

In replicate lineages that evolved on the spatially structured environment in the presence of the community, two distinct sets of genotypes emerged. Two replicate lineages had an *H-index* value of 0.75, but shared no mutations with the other three replicate lineages evolved under the same conditions. These two replicates harbored mutations in the *treC* gene, a gene involved in the trehalose degradation pathway, converting trehalose-6-phosphate into glucose and glucose-6-phophate. Mutations in this gene point to metabolic adaptation to the nutrients in the environment rather than the antibiotics.^62^ The other three replicates evolved in the spatially structured environment in the presence of the community also had an *H-index* of 0.75 for each pairwise comparison, but did not share any mutations with the other two replicate lineages. These three replicate lineages acquired mutations in the *traC, mpl* and *pfkA* genes. The *traC* gene mediates cell-cell contact during conjugation.^63^ ^64^ The *traC* mutation in both replicates is a synonymous mutation, suggesting these do not affect functionality, except perhaps through codon bias.^65^ The other genes, *mpl and pfkA* and play a role in cell wall biogenesis and carbohydrate degradation, respectively. The *mpL* gene contains a frameshift mutation, likely leading to a loss of function. The gene encodes UDP-N-acetylmuramate, part of the cell-wall recycling machinery.^66^ The *pfkA* gene harbored either missense mutations or an in-frame deletion. The gene encodes phosphofructokinase, and is part of the glycolysis pathway. *PfkA* is responsible for about 95% of phosphofructokinase activity in *E. coli.*^67^ Disruption of *pfkA* breaks the glycolysis pathway, redirecting fructose 6-phosphate toward alternative pathways such as the pentose phosphate pathway, which is important for the generation of NADPH.^68^ One gene that was mutated in all *E. coli* replicates evolved within the spatially structured environment in isolation was *nadR*, a component of the NAD salvage pathway.^69,70^ This gene converts β-nicotinamide D-ribonucleotide into diphosphate and NAD^+^. The mutations detected were either missense mutations or deletions, likely resulting in loss of function. Inhibition of *nadR* has been shown to increase cellular NAD/NADH levels. Because NADH is an important regulator for enzymes involved in metabolism, these mutations may therefore represent an adaptation to the nutrients conditions rather than to antibiotics.^71^ Additionally, elevated levels of NADH have been associated with efflux-mediated antibiotic resistance ^71^, suggesting a potential indirect contribution to resistance. Four out of five of the replicates also had mutations within the *adhE* gene, which encodes aldehyde-alcohol dehydrogenase.^72^ It catalyzes the reaction of acetaldehyde into acetyl-CoA, and also generates NADH. Together, the recurrent mutations in *nadR* and *adhE* point towards selection of NADH-associated metabolic pathways in spatially structured environments, potentially linking metabolic adaptation to antibiotic tolerance.

Overall, evolution within the spatially structured environments containing TMP-SMX was associated with mutations in genes, *nadR*, *adhE* and *pfkA,* predicted to affect cellular NADH/NADPH levels in *E. coli* in eight out of ten replicates. A previous study has shown that NADPH also contributes to the stabilization of the dihydrofolate reductase-ligand complex,^73^ with changes in this complex being signs of trimethoprim resistance. Interestingly, the eight replicates with mutations affecting NAD/NADPH-associated pathways co-occurred with elevated resistance gene copy numbers in lineages evolved in the spatially structured environment (Figure 3A). This suggests that mutations in *nadR, adhE* and *pfkA* may be co-adaptations correlated with increases in resistance gene copy numbers.

Additionally, mutations within prophages were detected in mutant *E. coli* chromosomes (Table S1). SNPs and complex insertions were observed predominantly in the spatially structured environments. We chose to exclude these from our H-index analysis because prophage regions often evolve at different rates compared to the core bacterial genome and undergo decay after integration^74^, and phage activity was not monitored during the evolutionary experiment.

We observed more evolutionary parallelism at the COG-level, consistent with previous research (Figure 4B).^54,75^ Most of the parallelism occurred within replicates evolved in spatially structured environments, both in the presence and absence of the community. In these groups, evolution within the spatially structured environments lead to both parallel genotypic changes (Figure 4) as well as parallel phenotypic adaptation (Figure 1AB). In contrast, lineages evolved in the well-mixed environment, in the presence of the community, showed parallel phenotypic adaptation without corresponding genomic parallelism, contrary to our expectation that stronger selection and larger population sizes would lead to more genetic parallelism.

In summary, we observed genetic adaptations at different levels of organization, including the integron cassette harboring resistance genes and mutations in the genomic background underly parallel phenotypic evolutionary trajectories of antibiotic resistance. However, the canalization of antibiotic resistance phenotypes could not be fully explained by genetic substitutions only, highlighting the role of additional factors, including the microbial community. These findings indicate that antibiotic resistance evolution is shaped by multiple levels of genetic of genetic and ecological organization.

## Conclusion and implications

We observed that the presence of a community has a protective effect on *E. coli* at the onset of the antibiotic resistance evolution experiment, an effect that diminished with increasing antibiotic concentrations. Community presence was associated with a faster evolutionary response in both the well-mixed liquid as well as spatially structured solid environments, consistent with previous findings showing that the presence of other bacteria can speed up and canalize evolutionary trajectories.^15^ Phenotypic parallelism was observed across all environmental contexts, either community or environmental structure, except in the well-mixed environment in the absence of the community. In spatially structured environments, the phenotypic parallelism was associated with genetic parallelism. In contrast, in the well-mixed environments, the phenotypic canalization in the presence of the community cannot solely be explained by genetic changes, indicating that non-genetic changes, such as phenotypic plasticity or physiological changes mediated by the community, likely contributed to the evolutionary response to antibiotics.

The beneficial effect of a community on antibiotic resistance evolution at sub-MIC levels has been observed before.^8,76^ Here, we show that the beneficial effect can persist after antibiotic concentrations exceed the initial MIC. Because the AUM environments used in this study are relatively nutrient poor, they likely provide opportunities for positive social interactions among microbes, which may contribute to the sustained beneficial effects observed.^77,78^ Theory suggests that communities benefit more from public goods interactions in structured environments, compared to well-mixed environments,^34,35,79,80–82^. However, during the experiment the positive effect of the community on *E.coli* on population size diminished with increasing antibiotic concentrations, over time (Figure 1CD), this was associated with smaller population sizes of the community members.

In the spatially structured environment, community presence did not have a large impact on the evolved MIC level of *E. coli*. However, the MBC increased when the *E. coli* lineages evolved in isolation were assessed in the presence of the community, indicating that the community-mediated protection remains relevant even for highly resistant lineages (Figure 2B). This effect was absent for *E. coli* evolved in the presence of the community in the well-mixed environment. These findings suggest that the beneficial effect of the community is not dependent on the antibiotic concentration alone. Instead, it depends on the evolutionary history of the *E. coli*, particularly whether lineages evolved in a community context. This highlights the importance of the ecological history in shaping antibiotic tolerance and resistance.

Initial cell density in antibiotic-containing environments is known to influence evolutionary outcomes.^17,76^ In both tested environments, *E. coli* reached higher densities when embedded within the community, which was associated with faster evolutionary trajectories towards high antibiotic resistance. This pattern is consistent with the well-known inoculum effect.^16,17^ However, for the *E. coli* evolved on the spatial structure in the presence of the community, this did not lead to a higher MIC compared to the *E. coli* evolved in isolation (Figure 1A,B).^8^ Because MICs were determined at equal *E.coli* population sizes, this assay does not take into account population size differences during the serial transfer experiments.^16^ Nevertheless, population size may indirectly contribute to reaching higher MICs, as the *E. coli* that evolved larger population sizes over 24 hours of growth in the absence of antibiotic reached higher MICs. This suggests that adaptation to the medium-environment contributes to the growth at higher antibiotic concentrations.

The high degree of copy number variation of the integron cassette carrying the antibiotic resistance genes, both within and across replicates, indicates substantial heterogeneity within replicate populations and suggests a dynamic process in which elevated copy numbers represent a potentially unstable state, likely maintained only under high selective pressure. This interpretation is in line with an insertion sequence-driven amplification mechanism at the resistance locus that can stochastically generate new copies under stress. At the same time, increased CNVs can lead to genomic instability by elevating the probability of disruptive insertions or deleterious homologous recombination events. Such a negative feedback mechanism could promote the continuous generation and purging of excess resistance loci within plasmid lineages.^83^ Previous studies have underscored the complexity of plasmid replication and maintenance, suggesting that various factors, including selective pressures and growth rates, can influence the stability of plasmid copy numbers. ^84,85,86^ The disconnect between resistance gene copy numbers and MICs in our experimental context echoes findings in such systems, where researchers have reported instances of decoupling between plasmid abundance and phenotypic resistance. ^87,88^ These observations challenge assumptions regarding the straightforward relationship between the presence of antibiotic resistance genes, gene expression and antibiotic resistance. In addition, the mobility of the resistance genes on the integron cassette may further contribute to variation and efficacy of the resistance genes. While our analysis allows us to determine gene copy numbers, it does not resolve the location of individual copies within the plasmid or genome, limiting our ability to predict their functional impact. Because gene expression is influenced by genomic context, including gene location, alignment with regulatory elements, and potential compensatory mutations such as those related to NADH observed in the spatially-structured environment, illustrate the challenges of relying solely on sequence-based approaches in the clinical detection of antimicrobial resistance.

Together, these findings demonstrate that repeatable antibiotic resistance evolution reflects the combined influence of multiple levels of genetic adaptation, environmental structure and ecological interactions and highlight the need for comprehensive investigations into the multifaceted ecological and genetic interactions governing plasmid-mediated resistance.

## Author contribution

Conceived and designed the study: LEZ MGJDV. Performed the experiments: LEZ MB JML. Analyzed the data: LEZ TH MGJ. Wrote the manuscript: LEZ JML TH MGJ.

## Acknowledgements

LEZ was supported by the Adaptive Life program of the Faculty of Science and Engineering, within GELIFES. MB was supported by the Faculty of Science and Engineering - Adaptive Life PhD Scholarship from the University of Groningen, awarded by GELIFES.

## Data

PRJEB90332

