## Supplementary Information for "When things add up: environmental structure and microbial interactions drive antibiotic resistance plasmid evolution"

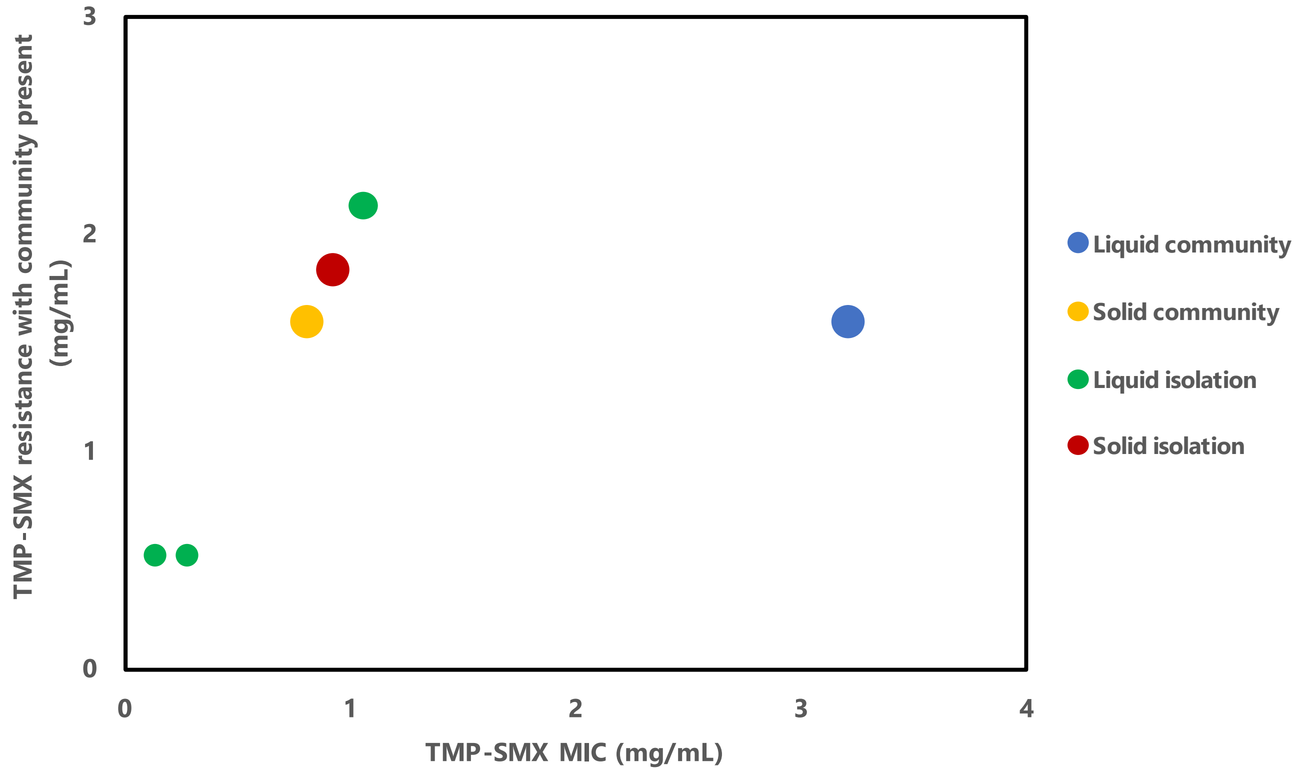

**Figure S1 | Minimal inhibitory concentration (MIC) of evolved *E. coli* in isolation correlated with antibiotic resistance of evolved *E. coli* in the presence of the community.** Concentrations of trimethoprim-sulfamethoxazole (1:5 ratio, TMP:SMX) are given by the trimethoprim concentrations. Size of the data points show the number of replicates, with the smallest data point being one replicate and the largest all five replicates. The community effect on MIC was determined by performing an MBC assay in the presence of the community. The minimum bactericidal concentration (MBC) of trimethoprim-sulfamethoxazole of *E. coli* on agar in presence of the community is depicted on the y-axis.

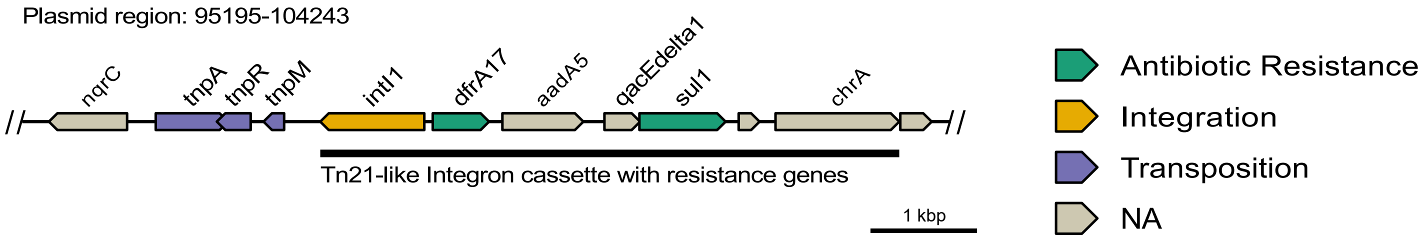

**Figure S2 | Structure and genes of the region within the plasmid conferring resistance to trimethoprim and sulfamethoxazole.** The *dfrA17* gene encodes a trimethoprim resistant dihydrofolate reductase protein and the *sul1* gene encodes a sulfonamide resistant dihydropteroate synthase protein, these are shown in green. Both resistance genes are located within a transposon, with the transposition-related genes being marked in purple. Of the transposition-related genes, *tnpA* encodes the IS*6* family transposase, *tnpR* a Tn*21* resolvase and *tnpM* encodes the Tn*21* modulator protein. The integration gene *intI1* that encodes a class 1 integron integrase, is shown in yellow. Other genes found within the integron cassette are *aadA5* which encodes aminoglycoside nucleotidyltransferase and confers resistance to aminoglycosides, *qacEdelta1* which encodes a quaternary ammonium compound efflux SMR transporter which confers resistance to a variety of ammonium compounds, and *chrA* which encodes a chromate transport protein which confers resistance to chromate.

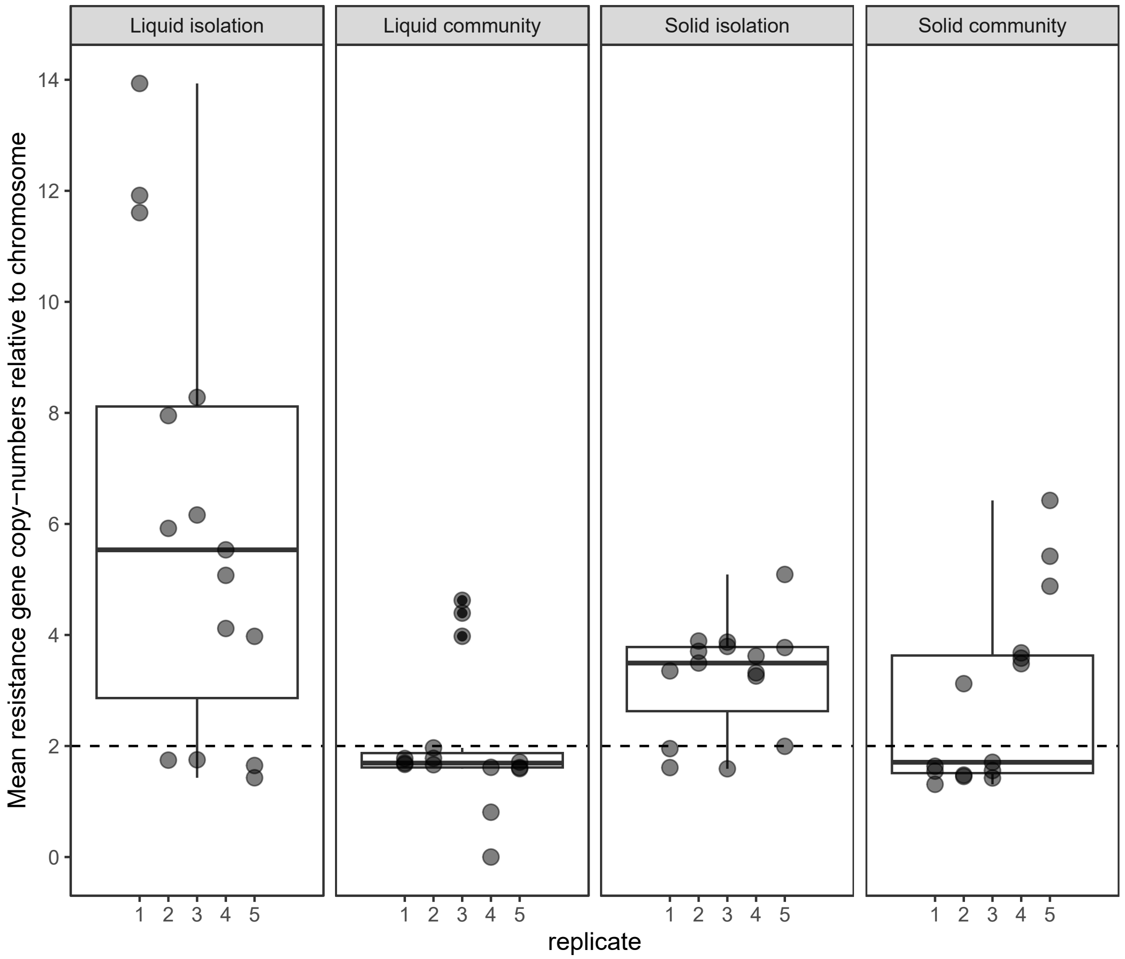

Liquid isolation
Liquid community
Solid isolation
Solid community
Mean resistance gene copy−numbers relative to chromosome
0
2
4
6
8
10
12
14
1 2 3 4 5
replicate

**Figure S3 | Copy number variations (CNVs) of the integron cassette containing antibiotic resistance genes.** Data shown are the CNVs after the 120 days of evolution in the final mutants. Three clones per replicate lineage were sequenced. The native isolate of *E. coli* contained a plasmid with the integron cassette shown in Figure S2. This cassette contains the genes *dfrA17* and *sul1* which confer resistance to trimethoprim and sulfamethoxazole respectively. CNVs are mostly increased when *E. coli* is evolved in isolation, but a few replicates of *E. coli* evolved in presence of the community showed increased CNVs. CNVs were significantly different for *E. coli* evolved in the well-mixed liquid environment in isolation and the three other environments ($P$=0.0030, $P$=0.025, $P$=0.024 for *E. coli* mutants evolved in the well-mixed environment with community present, and *E. coli* mutants evolved on the spatially structured solid environment in isolation and with the community present respectively, *Welch's t-tests*). There was also a significant difference for the CNVs of *E. coli* mutants evolved in the well-mixed environment with the community present and *E. coli* mutants evolved in the spatially structured environment in isolation ($P$=0.018), but no significant difference in CNVs between *E. coli* mutants evolved in the spatially structured environment with community present and those evolved in the well-mixed environment with community present ($P$=0.060), and those evolved in the spatially structured environment in isolation ($P$=0.87).

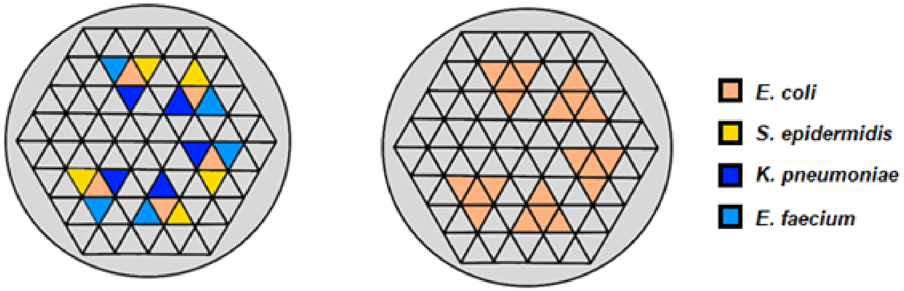

**Figure S4 | Setup of agar gel within 100 mm petri dish for spatially structured environments.** Left figure shows the community setup for all five replicates, right figure the control setup. For both environments, *E. coli* that is surrounded by either community members or other *E. coli,* was transferred. Bacteria were pipetted in the middle of each chamber on the AUM agar, and allowed to grow for 48 hours before the next transfer. We did not observe any spillover of colonies to other compartments.

**Table S1 | Type of mutations found on the *E. coli* chromosome of final propagations step mutants.** Each mutation is denoted with an “X”. Five replicate lineages per environment. Mutations are only shown if two of the three sequenced clones per replicate lineage contained that particular mutation. If a gene contained multiple unique mutations, they are shown on different rows.

Mutations in integrated prophage genomes are shown at the bottom of the table, with complex mutations indicating a combination of insertions and SNPs. The data from this table without prophage mutations were used to calculate the repeatability of evolution shown in Figure 4.

| Gene | Product | Mutation | Liquid isolation | | | | | Liquid community | | | | | Solid isolation | | | | | Solid community | | | | |
|---|---|---|---|---|---|---|---|---|---|---|---|---|---|---|---|---|---|---|---|---|---|---|
| | | | 1 | 2 | 3 | 4 | 5 | 1 | 2 | 3 | 4 | 5 | 1 | 2 | 3 | 4 | 5 | 1 | 2 | 3 | 4 | 5 |
| adhE | bifunctional acetaldehyde-CoA/alcohol dehydrogenase | snp | | | | | | | | | | | | | x | x | | | | | | |
| | | snp | | | | | | | | | | | | | | | x | | | | | |
| | | snp | | | | | | | | | | | x | | | | | | | | | |
| betT | Choline-glycine betaine transporter | snp | | x | | | | | | | | | | | | | | | | | | |
| btsR | two-component system response regulator | del | | | | | x | | | | | | | | | | | | | | | |
| btsS | two-component regulatory system sensor histidine kinase | snp | x | | | | | | | | | | | | | | | | | | | |
| cspE | transcription antiterminator/RNA stability regulator | snp | | | | | | | | | | | x | | | | | | | | | |
| cyaA | class I adenylate cyclase | snp | | | | | x | | | | | | | | | | | | | | | |
| folP | dihydropteroate synthase | del | | | | | | x | x | | | | | | | | | | | | | |
| ftsB | cell division protein | snp | | | | | | | | | | | | | | | | | | | | x |
| hrpA | ATP-dependent RNA helicase | snp | | | | | | | | | | | | | | x | | | | | | |
| iutA | ferric aerobactin receptor | snp | | | | | | | | | | | | | | | | x | | | | |
| kfiB | KfiB protein | del | | | | | | | | | | | | | | x | | | | | | |
| metH | methionine synthase | snp | | | | | | | | | | | | | x | x | | | | | | |
| mpl | UDP-N-acetylmuramate:L-alanyl-gamma-D-glutamyl-meso-diaminopimelate ligase | ins | | | | | | | | | | | | | | | | | x | | x | x |
| msyB | MysB family protein | snp | | | | | | | | | | | | x | | | | | | | | |
| mutY | A/G-specific adenine glycosylase | snp | | | x | | | | | | | | | | | | | | | | | |
| nadR | nicotinamide-nucleotide adenylyltransferase/ribosylnicotinamide kinase | del | | | | | | | | | | | x | x | | | | | | | | |
| | | snp | | | | | | | | | | | | | x | x | | | | | | |
| | | snp | | | | | | | | | | | | | | | x | | | | | |
| pfkA | 6-phosphofructokinase | snp | | | | | | | | | | | | | | | | | x | | | |

| | | | | | | | | | | | | | | | | | | | | | | |
|---|---|---|---|---|---|---|---|---|---|---|---|---|---|---|---|---|---|---|---|---|---|---|
| | | snp | | | | | | | | | | | | | | | | | | | | x |
| | | del | | | | | | | | | | | | | | | | | | | x | |
| radC | DNA repair protein | snp | | | | | | | | | | | | | | | | | | x | | |
| rbsR | ribose operon transcriptional repressor | del | x | | | | | | | | | | | | | | | | | | | |
| rpoC | DNA-directed RNA polymerase subunit beta' | snp | | | | | | | | | | | | | | | | | | | x | |
| | | ins | | | | | | | | x | | | | | | | | | | | | |
| rpoD | RNA polymerase sigma factor | snp | x | | | | | | | | | | | | | | | | | | | |
| sspA | stringent starvation protein A | snp | x | | | | | | | | | | | | | | | | | | | |
| tnp | IS3 family IS2 transposase ORF B | snp | | | | | | | | | | | x | | | | | | | | | |
| traC | type IV secretion system protein | snp | | | | | | | | | | | | | | | | | x | | x | x |
| treC | alpha,alpha-phosphotrehalase | snp | | | | | | | | | | | | | | | | x | | x | | |
| caiA | Carnitine operon oxidoreductase | snp | | | | | | | | | | x | | | | | | | | | | |
| yhcC | TIGR01212 family radical SAM protein | snp | | | | | | | | x | | | | | | | | | | | | |
| prophage 1 | phage tail | complex | | x | | | | | x | | | | x | x | x | x | x | x | | x | x | |
| | | snp | | x | | | | | x | | | | | x | | x | x | x | | x | x | |
| | | snp | | x | | | | | | | | | | x | | x | x | x | | x | x | |
| | | snp | | | | | | | | | | | | x | | x | x | | | | x | |
| prophage 1 | downstream tail assembly chaperone | complex | | | | | | | | | | | x | | | | | | | | | |
| | | snp | | | | | | | | | | | | | | | x | | | | | |
| | | snp | | | | | | | x | | | | | x | | | x | x | | x | x | |
| | | snp | | x | | | | | x | | | | | x | | x | x | x | | x | x | |
| | | complex | | x | | | | | x | | | | | x | | x | x | x | | x | x | |
| | | snp | | | | | | | | | | | | | | | x | | | | | |
| | | snp | | | | | | | | | | | | | | | x | | | | | |
| prophage 2 | downstream phage tail | snp | | | | | | | | | | | | | | | | | | | x | |

### Graphical abstract

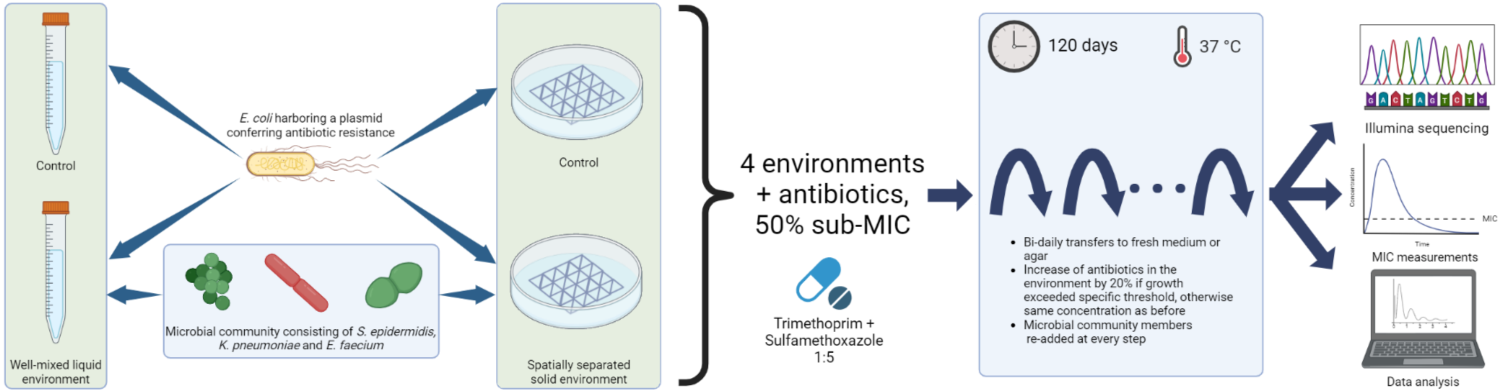
